# Context-Dependent Motor Feedback Underlies Choice-Related Signals in Visual Cortex

**DOI:** 10.64898/2026.01.05.697619

**Authors:** Nan Zhang, Yong Gu

## Abstract

Choice-related neural activity, reflecting trial-by-trial correlation with subjects’ perceptual choice, is commonly observed in sensory cortices and has been implied in sensory readout. Recent studies suggest an alternative source from motor-reporting, yet its precise nature remain elusive due to limitations in conventional experimental paradigms. Here we quantitatively identified motor component by training macaques to discriminate visual motion directions via making saccade to spatial-independent, colored choice targets appeared under two contexts: at stimulus onset, or after its offset. Neural recordings in visual areas MT/MST revealed a dominant motor-related choice signal when the saccade goal was known in advance; this signal vanished with delayed target onset. By comparing with downstream sensory-motor transformation areas and employing a recurrent network model, we demonstrated that this motor component reflected feedback. Our work clarifies that a major portion of “decision” activity in sensory neurons is attributable to motor-reporting processes, highlighting the profound influence of behavioral context.

## Introduction

Choice-related activity of single neurons has been frequently observed in many sensory systems, including visual^1–5^, somatosensory^6,7^, and vestibular pathway^8–11^. Specifically in a typical two-alternative forced choice (2-AFC) sensory discrimination task, it is found that some individual neurons exhibit significant correlations with the animals’ upcoming choice on a trial-by-trial basis. Amazingly, many neurons appear to be able to predict the animals’ choice with respect to their encoded task-relevant feature (the so called “choice probability”, or CP^2^. That is, the animals tend to make a choice towards the target associated with the sensory stimulus that is consistent with the neuron’s preferred feature, for example, motion direction of a patch of moving dots. Hence, it is ideal to infer a feedforward, sensory driven mechanism that mediates this process: sensory information carried by individual neurons is “read out” by downstream neurons, which consequently drives choice in downstream decision-related areas^12–14^, such as those sensory-motor association areas in the posterior parietal area (PPC), frontal and prefrontal cortex (PFC), superior colliculus (SC), striatum, etc.

However, a second, alternative hypothesis may also be possible: the observed choice-related activity in earlier sensory areas may reflect a top-down process in which a decision-related signal is sent back from higher level areas^15–19^. This hypothesis receives supports from a few sources of evidence. For example, psychophysical experiments have shown that subjects tend to rely more on sensory input during the early phase of the stimulus duration, yet choice-related activity is significantly delayed^18,20^. This mismatch in the temporal profile of the two metrics (psychophysical kernel vs. CP), is more consistent with the feedback than the feedforward process. By causal manipulations, recent studies demonstrate that choice-related activity in early visual cortex is largely diminished after inactivating the higher level areas^21^. A recent microstimulation experiment also reports that choice-related activity is not necessarily matching the microstimulation effect that indicates causality of sensory readout^22^. In summary, choice-related activity in early sensory areas could arise from a number of sources other than the bottom-up readout process, including prior knowledge^23–25^ in higher level areas developed during learning processes^26,27^, attentional modulation^15,28^, or correlated noise among single neurons^13,29,30^.

While previous research about choice-related activity is largely concentrated on feedforward vs. feedback, potential motor-related contributions have received little attention and are rarely investigated so far. In all perceptual-decision making tasks, the animals have to learn to correctly associate motor output to certain sensory input, a process shaped by reward reinforcement. For example, the animals need to make either saccadic eye movements, or hand reach to one of the choice targets displayed on the visual screen in front of the animals (e.g., left vs. right, or up vs. down). These motor-related signals are ubiquitously encoded in sensory-motor association areas (PPC, PFC, SC, striatum, etc.), and may affect choice-related activity in upstream sensory areas. Although there are few studies suggesting influence of motor-related source^17,31^, direct evidence identifying exact contributions from this source to previously observed choice-related activity, i.e., CP is rare, particularly in the extrastriate visual cortex, such as the middle temporal area (MT) and the medial superior temporal area (MST).

Choice-related activity, or CP, has been frequently reported in both MT and MST^1,2,8,32^. Yet in all these studies, CP has been measured by placing choice targets at fixed spatial locations, meaning a fixed relationship between the motor direction and sensory input (e.g., saccadic to the left target corresponds to left-moving random dots). In the current study, to quantify potential motor-related contributions to choice-related activity in MT and MST, we employed a colored-target paradigm^33,34^ in which monkeys were required to saccade to choice targets based on their color instead of fixed spatial locations, allowing dissociation of the two sources for choice-modulated signals: saccadic-related vs. perceptual-related. Through this paradigm, we provided solid evidence showing that in MT/MST, saccadic motor-related contribution dominates in shaping the CP. Importantly, this signal emerged later in MT/MST than in the downstream areas of the lateral intraparietal area (LIP), indicating feedback. All these results were reproduced by an end-to-end, two-module recurrent neural network (RNN). Importantly, manipulations in the RNN demonstrated that the feedback connections between modules are key to the saccadic choice signals.

## Result

### Behavioral performance

Two male rhesus monkeys were trained to discriminate visual motion directions of optic flow moving leftward or rightward, with task difficulty modulated by different coherence levels in the flow dots^35^. The animals reported their perceptual decisions by making saccadic eye movements to one of two colored targets (green or red) presented laterally on each side of the visual display. Target colors were consistently associated with motion directions, irrespective of their spatial locations (Figures 1A and 1B). Importantly, trials with colored targets on two different spatial locations (map1: green on left and red on right, map2: red on left and green on right) were randomly interleaved within each experimental session. This design decoupled the monkeys’ perceptual choice from motor response towards specific spatial locations. Furthermore, two versions of the experimental paradigms regarding the timing of emerging choice targets were designed: choice targets appeared as early as visual motion onset (early-target-onset, Figure 1A), or delayed until visual motion offset (delayed-target-onset, Figure 1B). The delayed-target-onset paradigm theoretically prevented motor planning during stimulus presentation, thereby it could serve as a control to eliminate potential motor effect as might be present in the early-target-onset condition. The early-target-onset paradigm was run first as a regular task in the current study, and when possible, the delayed-target-onset paradigm was further conducted in a subsequent block.

**Figure 1.**
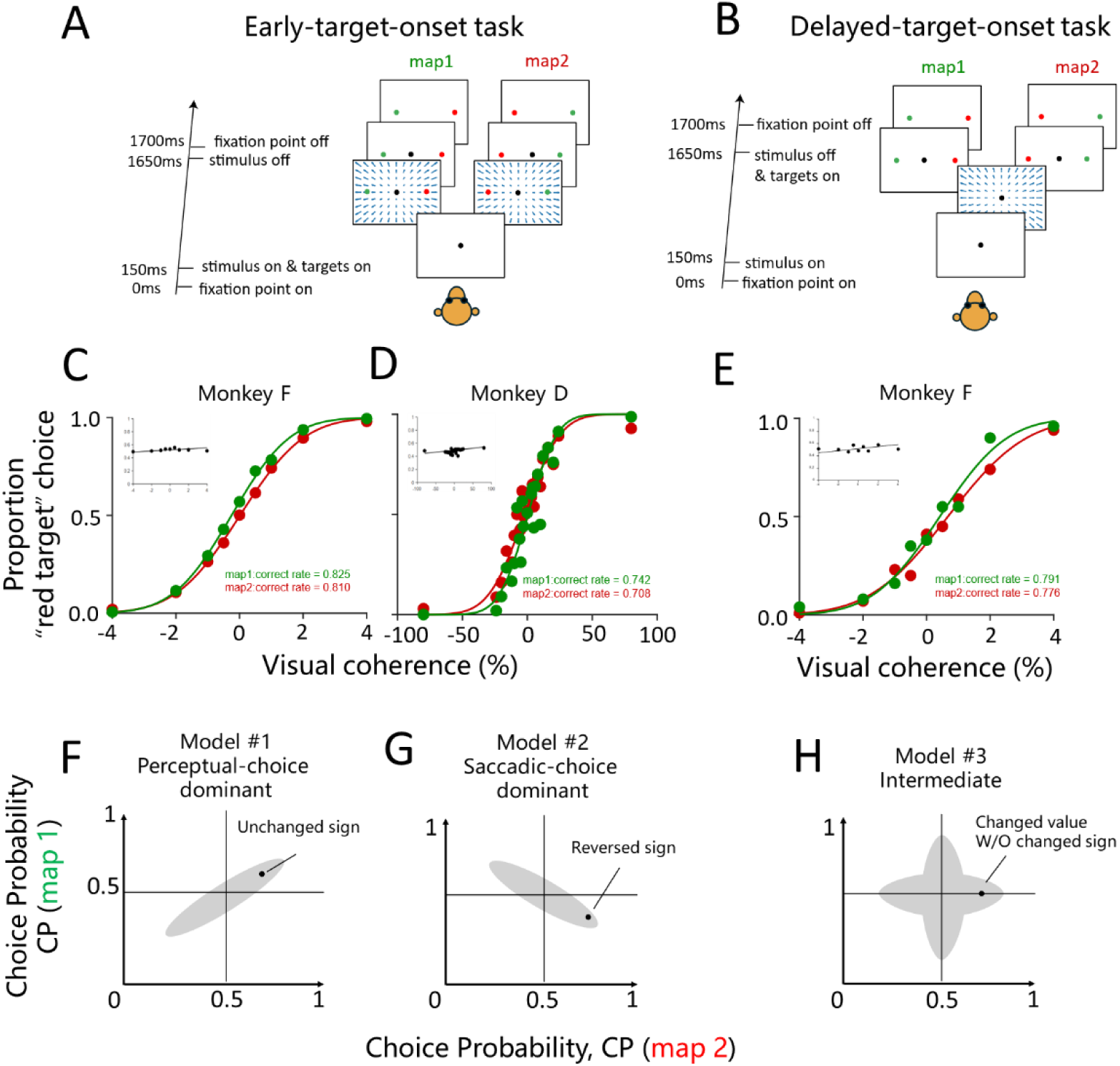
Disentangle perceptual and saccade choice using colored target task. (A) Task paradigm of early-target-onset. Two colored choice targets emerge simultaneously with the visual motion stimulus. Across trials, the green target could be either on the left (map 1), or on the right (map 2). (B) Task paradigm of delayed-target-onset. The colored choice targets do not emerge until the offset of the visual motion stimulus. (C-E) Behavioral performance quantified by psychometric functions fitted by accumulative Gaussian functions of the animals in the visual motion discrimination tasks (early-target-onset: monkey F & D; delayed-target-onset: monkey F). Green and red symbols indicate performance under map 1 and 2, respectively. Inset plots on the top left of each figure are performance based on spatial locations of choice target (left vs. right regardless of color). (F-H) Toy models of three possible outcomes of CP measured under map 1 and map 2 conditions. Gray areas represent distribution of CPs across single neurons. Black dot indicate an example.

After trained, both animals show stable behavioral performance by correctly associating visual motion directions with red-vs.-green targets (Figures 1C and 1D), regardless of their spatial locations (Figures 1C and 1D, inset plot). One of the monkeys was able to further run the delayed-target-onset task by performing based on colors (Figure 1E) instead of spatial locations (Figure 1E, inset plot). We then performed single-unit recordings in a number of areas while the animals performed the tasks at the same time. In particular, we recorded from two visual motion areas in the superior temporal sulcus (STS): the medial superior temporal area (MST) and middle temporal area (MT). In addition, their downstream area in the intraparietal sulcus was also recorded: the lateral intraparietal area (LIP) (Figure S1). Overall, we found data were largely similar between areas MT and MST in the STS (Figure S2), while the difference was notably much larger between the two sulci (STS vs. IPS). Thus, in the following analyses, we grouped data between MT and MST to focus on comparison between STS and IPS.

### Disentangle perceptual and saccadic choice signals

Before we examine the neurophysiological data, we predict two possible, distinct CP patterns that exhibit opposite trend particularly from the early-target-onset experimental paradigm. First, if neural activity associated with choice reflects more abstract, perceptual decisions that are independent on saccadic spatial locations, the CP values would remain the same regardless of specific spatial locations of the choice targets. In this case, CP values on a cell-by-cell basis would be same between the two conditions of the spatial locations of the color targets, and the data would be along the positive unity line on a scatter plot (Model #1, Figure 1F). Alternatively, if choice-related activity mainly reflects saccadic spatial locations, the CP values would change by reversing their signs (> or < than the chance level of 0.5), depending on whether the choice is made to the left or to the right spatial target (Model #2, Figure 1G). Notably, a third hybrid model is also possible in which both factors contribute to the final choice-related signals, leading to changed CP value (but without having to change the sign), between the two target-mapping conditions (Model #3, Figure 1H). Nevertheless, our experimental paradigm allows a quantitative dissection and evaluation of each component.

In the neurophysiological data, we found both types of choice signals under the early-target-onset paradigm. For example, a neuron in the STS shows similar CP value between the two map conditions (green-on-left vs. red-on-left) throughout stimulus duration (Figure 2A, left panel, green vs. red symbols), indicating perceptual choice that was regardless of the targets’ spatial locations. Indeed, under the delayed-target-onset paradigm in which saccadic choice signals were unlikely to be present before presentation of choice targets on the visual display, the CP signals largely remained akin to those in the early-target-onset paradigm (Figure 2A, right panel). On the contrary, another exemplary STS neuron shows significantly changed CP value and its sign (> or < than the chance level of 0.5) between the two map conditions during the stimulus period in the early-target-onset task (Figure 2B, left panel), indicating saccadic choice that was highly dependent on the targets’ spatial locations. This motor-related effect was further confirmed by its disappearance in the delayed-target-onset paradigm (Figure 2B, right panel). In addition to STS, it is also frequent to see both perceptual-dominant (Figure 2C), and saccadic-dominant cells (Figure 2D) in the IPS areas.

**Figure 2.**
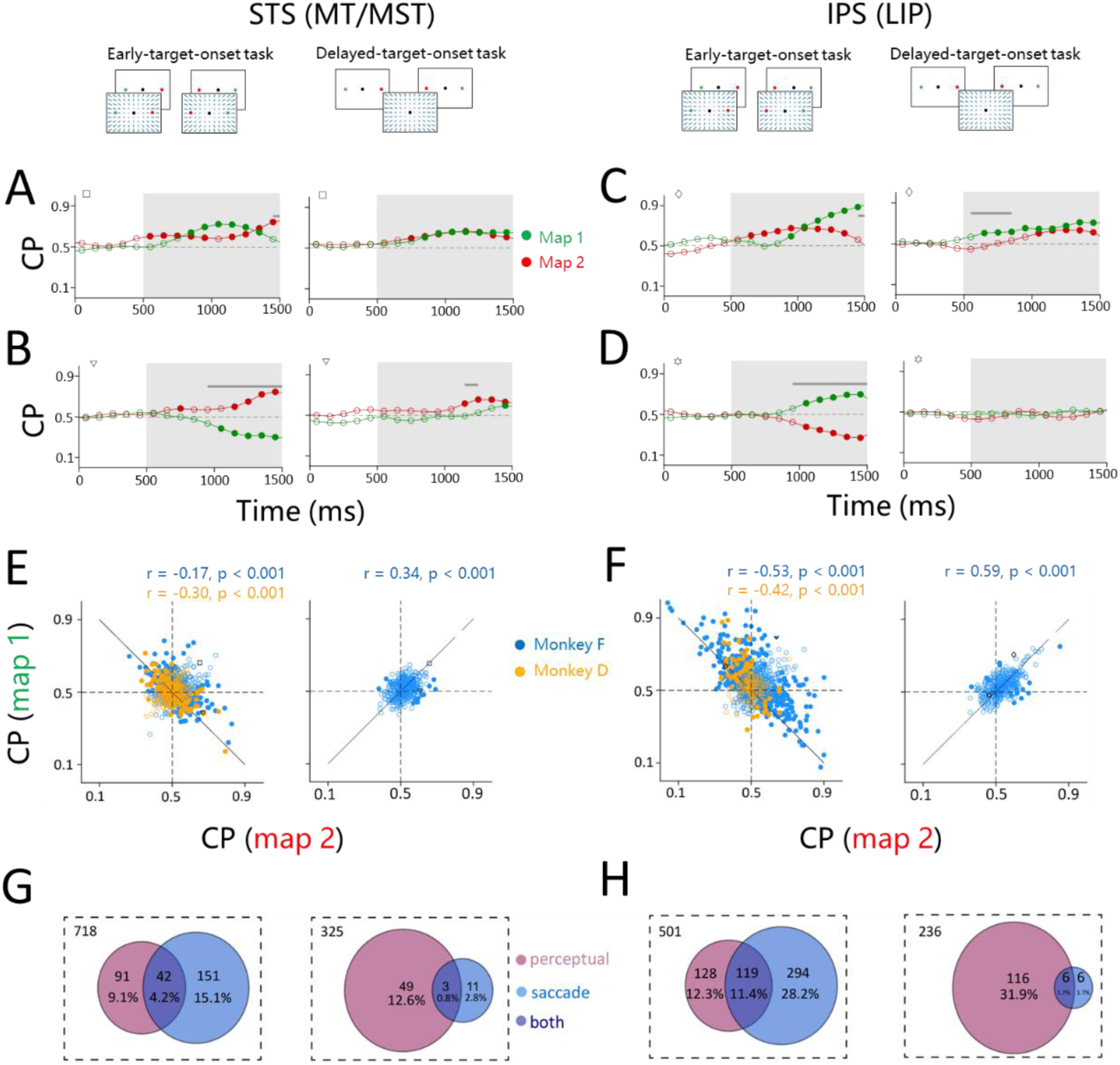
Quantify saccadic and perceptual choice component in CP. (A-D) Four exemplary neurons showing perceptual choice signal in STS (A) & IPS (C), and saccadic choice signal in STS (B) & IPS (D). CP measured under the early-target-onset and delayed-target-onset tasks are on the left and right in each plot, respectively. Green and red symbols represent the two maps. Horizontal gray bars indicate significant difference in the CP between the two maps. Filled and open dots are CP significantly and insignificantly different from the chance level (0.5), respectively. Gray areas indicate time window used to compute a single CP value for each neuron in the following analysis. Symbols on the upper left corner in each plot represent the exemplary neuron in the following population data (E and F). (E and F) Comparison of individual neuron’s CP under the two-map conditions in STS (E) and IPS (F) CP measured under the early-target-onset and delayed-target-onset tasks are on the left and right in each plot, respectively. Filled and open dots represent neurons with significant and insignificant saccadic choice component, respectively. Embedded texts inform Pearson correlation of CP measured between the two maps for each animal (blue, monkey F; orange, monkey D). Solid line is unity line. Horizontal and vertical dashed lines are chance level of CP (0.5). (G and H) Proportion of neurons encoding perceptual (magenta), saccadic (cyan), or both (purple) choice component in STS (G) and IPS (H). CP measured under the early-target-onset and delayed-target-onset tasks are on the left and right in each plot, respectively. Dashed rectangle indicates number of all sampled neurons.

We then computed a single CP value for each neuron under each choice-target location condition (map 1 vs. map 2), based on firing rate between 500 ms and 1500 ms, during which neural activity exhibited largest modulations (Figures 2A, 2B, 2C and 2D, shaded area). The CP value was then compared across population on a cell-by-cell basis between the two map conditions (green-on-left vs. red-on-left, Figures 2E and 2F). In the early-target-onset experimental paradigm, we found a significant negative correlation of CP between the two target locations in both STS (monkey D: r = −0.3, p < 0.001; monkey F: r = −0.17, p < 0.001, Pearson correlation, Figure 2E, left panel) and IPS (monkey D: r = −0.42, p < 0.001; monkey F: r = −0.53, p < 0.001, Pearson correlation, Figure 2F, left panel). Such a pattern is more consistent with the expectation model #2 (Figure 1G), indicating clear motor-related effect on the CP value. Indeed in the controlled delayed-target-onset paradigm, this negative correlation was vanished, and the pattern was reversed to show positive correlation in both STS (monkey F: r = 0.34, p < 0.001, Pearson correlation, Figure 2E, right panel) and IPS (monkey F: r = 0.59, p < 0.001, Pearson correlation, Figure 2F, right panel), meaning more consistent with the expectation model #1 (Figure 1F). This conversion confirms that overall saccadic choice signals are predominant over perceptual choice signals in the early-target-onset paradigm in both STS and IPS areas.

As can be seen in the population data (Figures 2E and 2F), some neurons in fact may carry both types of choice-modulated signal. To quantify exact contribution of each component, we computed perceptual-related and saccadic-related choice-modulated signals for each neuron. In particular, perceptual-related signal was quantified through ROC analysis with respect to color choice, irrespective of their spatial locations. On contrary, saccadic-related signal was quantified through ROC analysis based on spatial locations of the choice targets (left vs. right) instead of color (see Method). Significance of each signal was assessed by a re-sampling statistical test procedure (p < 0.05, permutation test). Across population, we then categorized those neurons with significant choice-modulated signals into one of the three categories: perceptual-only neuron, saccadic-only neuron, and mixed neurons (Figures 2G and 2H). It can be seen that under the early-target-onset experimental paradigm, a larger proportion of neurons encode saccadic choice signals than the perceptual ones in both STS (19.3% vs. 13.3%, Figure 2G, left panel) and IPS (39.6% vs. 23.7%, Figure 2H, left panel). Importantly, this pattern was reversed in the delayed-target-onset paradigm: far fewer neurons now showed saccadic choice signals in both STS (3.6%) and IPS (3.4%) (Figures 2G and 2H, right panels), supporting that a large proportion of motor-related choice signals were present during the early-target-onset experimental paradigm.

### Temporal dynamics of saccadic and perceptual choice-modulated activity

We then compared temporal evolution between the two types of signal (saccadic vs. perceptual) in the two sulci (STS vs. IPS). The goal was to infer possible feedforward vs. feedback information flow of each signal component.

To address this, we first simply averaged perceptual and saccadic choice-modulated activity across those neurons with significant modulations of each signal (Figures 2G and 2H) over the stimulus duration time in each brain region (Figure 3). This population analysis revealed that both animals gradually developed saccadic choice-modulated signal in the early-target-onset paradigm in both sulci (Figure 3A and 3B), while this signal was largely absent in the delayed-target-onset paradigm (Figure 3C). For perceptual choice, the signals also gradually developed in both areas under both early-target-onset (Figure 3D and 3E) and delayed-target-onset (Figure 3F) paradigm. Overall, the rising speed of the perceptual signals appeared to be faster than the saccadic ones. However, this analysis relied heavily on the criteria used for including the neurons in the dataset, which might introduce bias. For example, only those neurons with significant modulation (p < 0.05) based on activity between the 500 ms and 1500 ms time window after the stimulus onset were employed for the analysis.

**Figure 3.**
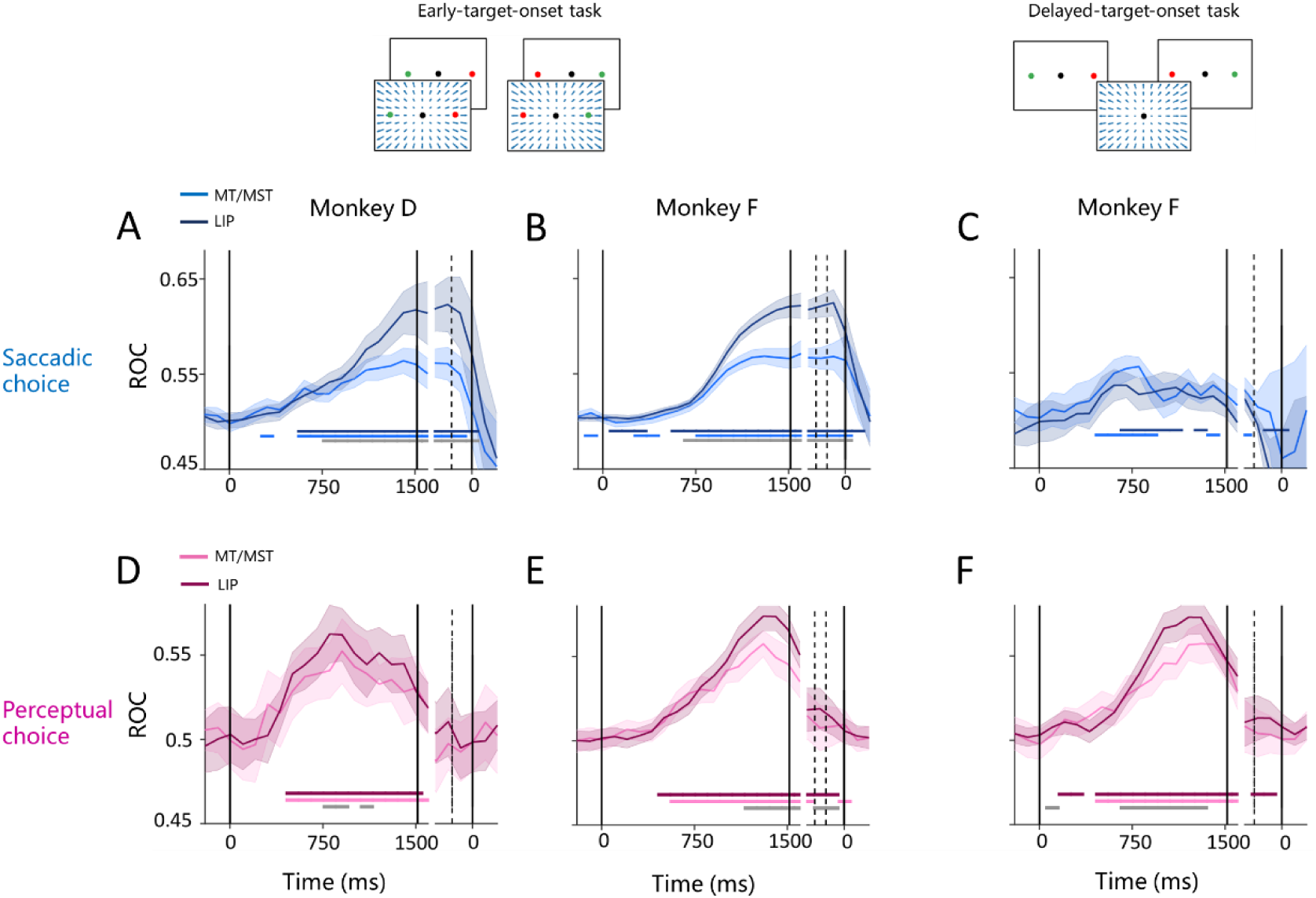
Temporal dynamics of perceptual and saccadic choice signal. (A-C) Temporal evolution of saccadic choice signal averaged across all those neurons with significant saccadic component as defined in Figures 2G and 2H. Dark blue: LIP; light blue: STS. Shaded areas 95%CI. Horizontal dark and light blue bars indicate significance from chance level. Gray horizontal bars indicate significant difference between the two areas. Data were shown for three animals collected under the early-target-onset (A and B) and delayed-target-onset paradigm (C). (D-F) Similar format in A-C for perceptual choice signal. Note that in contrast to the saccadic component that is largely gone in the delayed-target-onset condition (C), the perceptual component remains (F).

In order to better compare temporal evolution of the two choice signals across sulci, we further performed an analysis that was much released from restrictions that have been applied in the above analysis (i.e., Figure 3). In particular, we applied a model-based targeted dimension reduction (mTDR)^36^ based on all neurons on a time-to-time point across the whole stimulus duration (see Method). Through this analysis, both saccadic and perceptual choice signals were successfully captured and projected onto the first sequential principle component (see Method) of each relative subspace, allowing for comparison across areas. In both monkeys, it was shown that saccadic choice signals emerged significantly earlier in LIP than STS, with a medium lag of 60 ms in both monkey F (p < 0.001, rank-sum test, Figure 4A) and monkey D (p <0.001, rank-sum test, Figure 4B). On the contrary, perceptual choice signals emerged significantly earlier in STS than IPS with a medium lag of 20 ms in monkey F (p <0.001, rank-sum test, Figure 4C) and medium lag of 100 ms in monkey D (p <0.001, rank-sum test, Figure 4D). Importantly, in both areas, perceptual choice signal emerged earlier than the saccadic one (p < 0.001, rank-sum test). Therefore, these results indicated that perceptual choice signals might reflect feedforward information transmitted from STS to IPS, while the saccadic choice signals might mainly reflect feedback projections from IPS to STS. Interestingly, as revealed in the neural subspace, the saccadic choice signal is orthogonal to both of the perceptual and stimulus axis (Figure S3), indicating that it may exert limited influence on the latter two signals (see Discussion).

**Figure 4.**
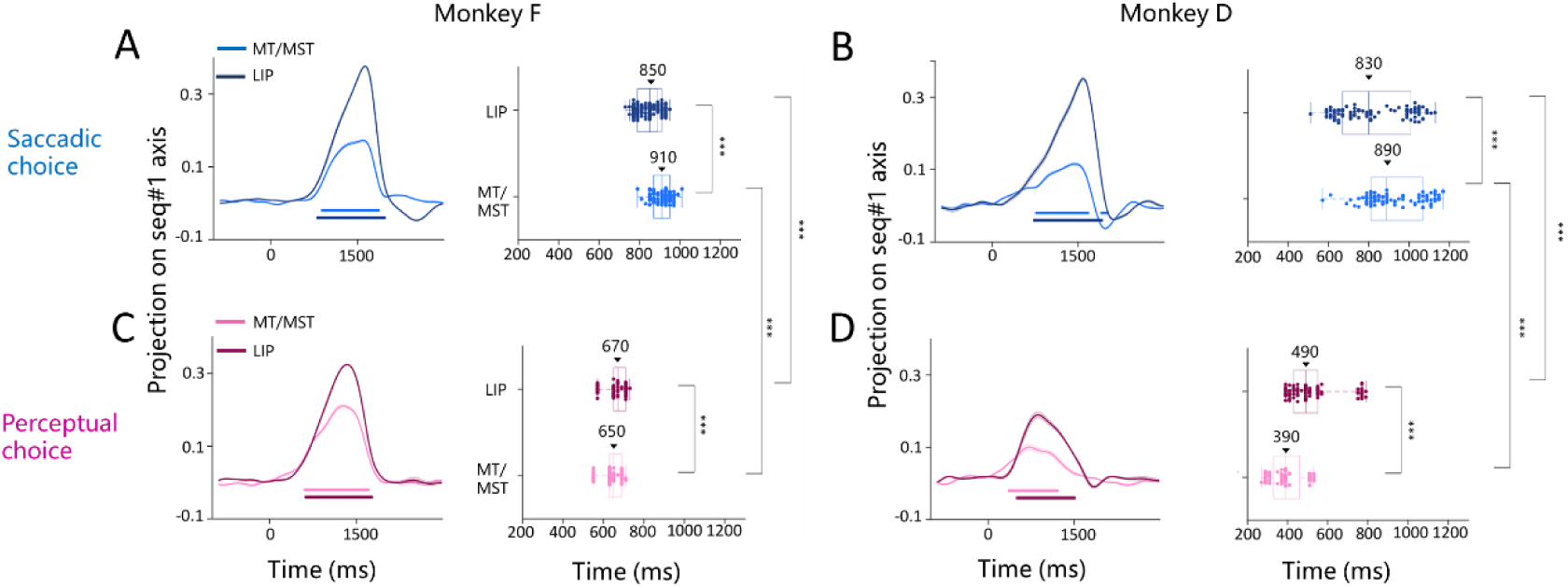
Quantify temporal delay between areas using PCA. (A and B) Comparison of temporal delay of saccadic choice component between STS (light blue) and LIP (dark blue) in each animal (A: monkey F; B: monkey D). In the left panels of each plot, population saccadic choice divergence signal was projected onto sequential principle component axis. Shaded bar represent 95%CI, by 100 times of resampling. In the right panel of each plot, each dot on the x-axis represents in each resampling, the time when saccadic choice signal is significantly larger than the base line. Number of asterisk indicate significant level of difference between the two areas: * p < 0.05; ** p < 0.01; *** p<0.001. (C and D). Similar format for perceptual component. Comparison was also made between the two types of signals (A vs. C; B vs. D).

### RNN simulations identify sources of motor and perceptual choice signal

To help understand the nature of saccadic-motor and perceptual related choice signals across areas, we constructed a two-module recurrent neural network (RNN). Specifically, module 1 receives visual motion input and processes directional information to simulate MT/MST in the STS. Module 2 receives visual motion information feedforwarded from module 1, as well as color information from the two choice targets, simulating sensory-motor transformation areas of LIP in the IPS. Importantly, module 2 also sends feedback signals to module 1 (Figure 5A), thus the connection between the two modules is bidirectional, consistent with previous anatomical findings^37^. The network’s goal was to correctly identify visual motion direction: either towards leftward or rightward. After training, the network reached satisfactory performance, in a way similar to the animals (Figure 5B), indicating that the RNN successfully learned the association between the flow direction and target color.

**Figure 5.**
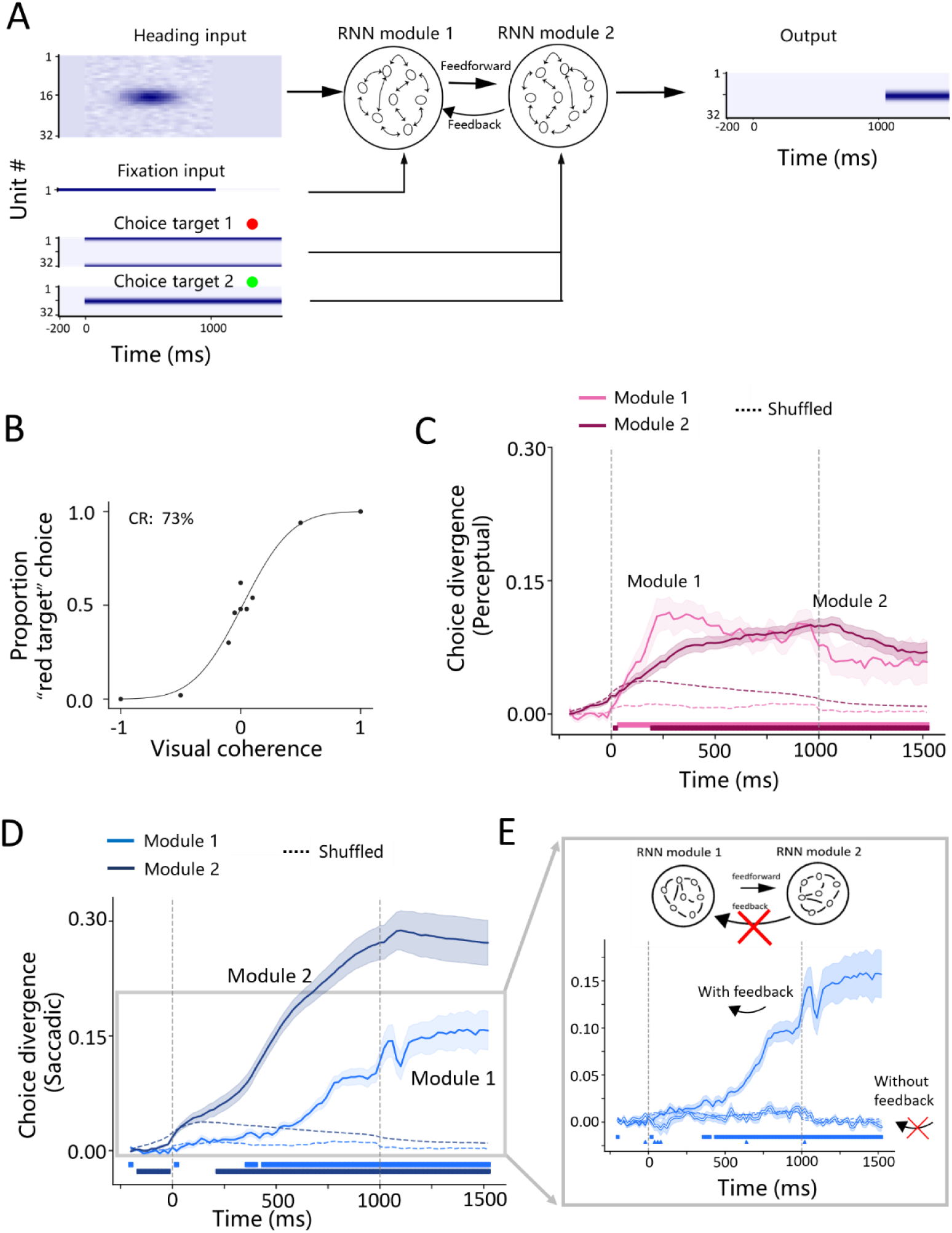
A two-module RNN simulation and manipulations reproduced saccadic and perceptual information flow across areas. (A) Construction of a two-module RNN simulating brain regions in STS and LIP. At the input level, the heat map represents response for the input module units in an exemplary trial, with darker color indicating higher firing rate. Input units were ranked according to their tuning directions (upper left). One additional unit encodes fixation point (middle left). These units were connected to the first module (RNN module 1). In addition, 32 units encode the red and green color targets, and they were connected to the second module (RNN module 2, bottom right). At the output level, 32 units encode the output decisions. There are 128 units in module 1 and 256 units in module 2. The units in each module are recurrently connected. Between modules, there are feedforward and feedback connections. All the connections could be manipulated. (B) Performance of an example network quantified by psychometrical function. (C) Temporal evolution of perceptual choice component between module 1 (Solid, light magenta curve) and module 2 (Solid, dark magenta curve). The data were averaged from 16 randomly initiated networks. Shaded area: 1SEM. Dashed curves: shuffled data. Horizontal bars: significant choice divergence from chance level. (D) Same as C but for saccadic choice divergence signal. (E) Same as the data in the gray area of E but with manipulation of the RNN feedback connections from module 2 to module 1 (e.g., remove, red cross). Squares and triangles represent significant saccadic signals before and after removing the feedback, respectively.

We then analyzed perceptual and saccadic choice-related signals that are possibly represented by the network’s hidden units in module 1 and module 2. Indeed we found both types of signals existing in both modules. To examine what factors may determine characteristics of these signals, we systematically manipulated connections between modules (feedforward vs. feedback).

Firstly for perceptual choice, we found that the signals tended to emerge earlier in module 1 than module 2 averaged from 16 networks in which initial weights were randomly assigned before training (Figure 5C).

Secondly for saccadic choice, we found a contrary pattern. That was, the signals tended to emerge earlier in module 2 than module 1 (Figure 5D). This pattern suggested that the saccadic signals might reflect feedback projection from module 2 to module 1. To verify this rationale, we applied two control manipulations. First, the feedback connection was removed in the network. It was shown that after cutting off this connection, the saccadic choice-related signal largely disappeared in module 1 (Figure 5F). Second, we rewired the choice target inputs to module 1 instead of module 2, meaning that the sensory signals could be directly transformed into motor choice in the module 1, rather than having to go through the higher level of module 2. In this case, we found that the saccadic choice in module 1 tended to emerge even earlier than the case when the choice target inputs were wired to module 2 (Figure S4). These manipulation results indicated that the onset of saccadic choice signals were highly dependent on where motor-related information is implemented.

## Discussion

Employing a colored choice-target paradigm, we were able to identify a motor-related (saccadic) component in choice-related activity within the sensory cortices of MT/MST. We also demonstrated that this signal dominated over perceptual-related component when measured in the early-target-onset context. In addition, the saccadic-related signal in MT/MST emerged later than that in the downstream areas of LIP, suggesting that this type of signal may arise from feedback projection from higher-level sensory-motor transformation areas. In contrast, motor-independent (perceptual) signals appeared earlier in MT/MST than LIP, more consistent with a feedforward processing. An artificial, two module RNN simulation largely reproduced the neurophysiological findings. Critically, network manipulations revealed that saccade-related motor choice signals depended on feedback connections between modules, whereas perceptual component depends on both feedforward and feedback connections. Thus, our study provides direct evidence that behavioral report (saccadic eye movements) exerts a strong motor influence on choice-related activity in sensory cortex. By dissociating and comparing motor-related and perceptual components in early sensory areas, we also identify their respective arising source.

### Origin of motor-related activity in the perceptual-decision circuit

Choice-modulated signals have been broadly observed in the perceptual-decision circuit, from low or mid-stage sensory areas to higher level sensory-motor association/transformation areas. In sensory-motor transformation areas, choice related signals are typically reflected as a ramping pattern over the stimulus presentation time, a process proposed as sensory evidence accumulation, and modeled most often as a drift diffusion process for decision making^38^. This choice modulated activity should be at least partly related to motor because the ramping direction of the neural activity (up vs. down) is tightly dependent on whether the upcoming choice is made toward or away from the response field of the neurons. In contrast, neural activities in lower level sensory areas typically do not exhibit a ramping pattern, yet they can also be modulated by choice as quantified by CP^2,15^. Previous debate focuses on whether CP in early sensory areas reflects a bottom-up, or a top-down information process, without much investigations into potential contributions from motor effect due to behavioral report. Recent studies, however, begin to notice this. For example, one recent study excludes motor-related choice component in the ventral visual pathway (V2/V3A) by using colored-target paradigm^17^, yet it remains elusive about the precise nature of this motor-related activity when compared to the perceptual component. In the dorsal visual pathway, a recent study discovers that CP of MT neurons are related with the relationship of their preferred motion direction and their receptive location, providing hints of existing motor-related influence in CP^31^. However, a quantitative identification and its comparison with perceptual component is infeasible due to choice targets have been placed in specific spatial locations with a fixed rule with respect to the sensory stimulus, leading to entangled motor-dependent (e.g., saccadic eye movement) and motor-independent (perceptual, or abstract) signals. In the current study, through a colored-choice-target experimental paradigm, we successfully dissected the two types of signals, and demonstrated surprisingly a strong motor-related component that dominated over the perceptual component. This immediately raises an important question: whether this motor related choice signal arises from early sensory areas, or it is a feedback from higher level sensory-motor transformation areas?

Both possibilities may exist. First for the early-motor-arising hypothesis, there is evidence suggesting that motor may be more tightly linked to earlier sensory areas than conventional impressions about their role in “pure” sensory processing. For example, a direct anatomical projection exists from MT to saccade control regions^39,40^. In another study, inactivating MT leads to choice bias towards the ipsilateral visual field, which is away from the receptive fields of the inactivated neurons, indicating some motor effects in MT^31^. In the MST, although saccadic responses are usually absent, many neurons are clearly modulated by smooth pursuit eye movements even in darkness^41,42^, indicating motor signals. It would be interesting to examine in a task when choice is made by smooth pursuit, whether a motor-related choice signal would directly arise from these areas. Finally, some studies have reported lack of perceptual signals in sensory-motor association areas including LIP^43^, PFC^44^ or PMd^45^, suggesting that sensory signals may not have to be transmitted to higher level areas, but may have been transformed into motor output decisions in the earlier sensory areas. In our RNN simulation, we find that directly wiring the color-target input to module 1 instead of module 2 results in earlier generation of saccadic choice signals in the former than in the latter (Figures S4A and S4B). Although there is rare evidence suggesting that MT/MST receives color information^46^, it would be interesting to examine whether saccadic choice signals may emerge earlier in the ventral visual pathway (where color information is encoded) than in the sensory-motor association areas.

Secondly, for the late-motor-arising hypothesis, motor-independent sensory information needs to be transmitted to higher level sensory motor association areas for evidence accumulation, and subsequent motor transformation. Supporting this, a few studies have reported abstract (perceptual) choice signal in sensory-motor association areas including LIP^47–49^, PFC^50,51^, SC^52^. A recent study in the prefrontal cortex (PFC) reported early rising of perceptual-related choice signals in a neural state subspace along an axis orthogonal to the axis encoding saccadic-choice signals^50^. This mechanism may even hold under conditions when choice targets are spatially anchored (instead of colored-target). For example, in a task when the monkeys are trained to discriminate visual motion directions by associating sensory stimulus to spatial targets with a fixed rule (e.g., left motion associated with left target, and vice versa for right motion with right target), it is suggested that the perceptual learning process happens in the sensory-motor association area of LIP instead of the upstream sensory area of MT^27^.

In summary, our current findings are more consistent with the second, late-motor-arising model, yet this may not be a universal rule in the brain. It is likely that in other areas (e.g., ventral visual pathway), or other behavioral report contexts (e.g., by smooth pursuit, or hand reach), the motor effect on choice-related signals may emerge earlier. Nevertheless, we predict that in whichever cases, a strong motor effect due to behavioral report would be present, which warrants future investigations.

### Functional implications of feedback saccadic signals

Choice-related signals in early to mid-stage sensory areas have long been interpreted as reflecting perceptual decisions. One piece of evidence supporting this rationale is that CP values are positively correlated with the neuronal sensitivity^1,2,4,8,12,53,54^, seemingly implying that more sensitive neurons tend to contribute more to the decision process. Yet there is also much evidence suggesting misalignment between the decisional choice and encoded sensory information (stimulus preference) ^55–57^. For example, using partial correlation analysis^55^, or dimensional reduction analysis^56^, it is found that choice related signal is largely orthogonal to the sensory axis in the extrastriate visual cortex. In our current study, we could further dissociate the choice signal into two components: saccadic-related and perceptual-related signal. And we find that In STS the saccadic choice axis is orthogonal to both of the stimulus axis (∼90 °) and the perceptual choice axis, while the latter two axes are not orthogonal to each other (∼60°) (Figure S3). This finding indicates that perceptual choice signal is related to sensory encoding, while the feedback saccadic choice signal is unlikely to interfere with sensory coding or the perceptual choice.

Numerous previous studies have reported feedback source of choice related signals in sensory areas, which may be linked to feature attention^15,29^, or priors^23,30^. A recent study using electrical microstimulation indicates that those choice-related signals inconsistent with sensory encoding are independent on the perturbation effect, suggesting a feedback rather than a feedfoward, sensor-driven source^22^. When dissociating choice-related activity into the motor-independent signal, a recent study demonstrates that even the perceptual-related choice signal in V2/V3A can arise from feedback^17^. In some of these previous studies, feedback choice signals may be ecologically relevant, yet the exact functional implications of the feedback saccadic signals remain elusive. One possibility is that the motor-modulated activity (e.g., due to saccade) may help the animal to be alert to spatial locations around the intended saccadic target. This is biologically plausible as it has been shown that sensory area activities are gained up by preceding saccadic command^58^, reflecting spatial attention into the sensory neurons’ receptive field. Alternatively, the feedback saccadic choice signal may be inevitably present in early sensory areas simply due to rich recurrent and feedback connections in the brain. Indeed, in our RNN simulation, cutting off the feedback connects from the higher level module to the lower level one abolished the motor related choice signal in the lower level module (Figures 5D and 5E). At the same time, the network’s behavioral performance remains unchanged (Figure S4C). Therefore, these motor-related signals may not have to exert significant influence on the circuit’s functions.

## Method

### Subject and surgery

Two male rhesus monkey (Macaca mulatta weighing 6∼9 kg) were used in our data collection. Before the behavioral training, the animals were chronically implanted with a circular molded lightweight plastic ring (containing a bottom ring, a middle ring and a lid). The plastic rings were served as both head-fixed post and neural recording chamber. After recovery, monkeys were trained to seat in a customized primate chair with head restraint. All animal procedures were approved by the Animal Care Committee of the Center for Excellence in Brain Science and Intelligence Technology, Chinese Academy of Sciences.

### Apparatus

The visual stimuli used in the current experiment were generated by OPENGL and controlled by Temponet software developed by Reflective Computing Inc. (USA). Random dots were presented on a 43-inch HDR1000 display screen. Eye movements of the animals were collected using an Eyelink OptiPlex 5040 eye tracker from SR Research (Canada), which tracks eye movements via infrared reflection signals. The water delivery system consisted of a Lange peristaltic pump controllable via digital signals to regulate reward duration. During the experiments, the experimental room was kept in dark except the residual light from the visual display. Viewing distance was 30 cm. The visual display was 50 cm in height and 60 cm in width, corresponding to visual angle ranges of approximately 90 by 90 degrees. Random dots were full of the screen to simulate self-motion through natural environment (optic flow). Motion parallax and size of dots information was provided to simulate depth. No horizontal disparity information was provided.

### Behavioral task

The animals were trained to perform a heading discrimination task based on the direction of the flow field relative to straight ahead (leftward vs. rightward). To report, the animals made saccadic eye movement to one of the two choice targets placed at the horizontal meridian. The two choice targets are green and red in color, corresponding to leftward and rightward heading direction, respectively. In each experimental block, the green target was placed on the left for 50% trials, and on the right for the rest trials. The two conditions were randomly interleaved on a trial by trial basis. There were two experimental paradigms regarding the time of the target emergence. In the early-target-onset condition, the choice targets were presented simultaneously with the optic flow. In the delayed-target-onset condition, the choice targets were not presented until the offset of the optic flow. These two conditions were run in different blocks. Task difficulty was controlled by visual coherence (monkey F: ±4%, ±2%, ±1%, ±0.5%, 0% for Early target onset task; ±4%, ±3.2%, ±2%, ±1.6%, ±1%, ±0.8%, ±0.5%, ±0.4%,0% for Delayed-target-onset task; monkey D: ±80%, ±24%, ±20%, ±16%, ±12%, ±10%, ±8%, ±5%, ±5%, ±4%, ±3%, 0% for Early-target-onset task). To simulate heading, visual motion was conducted using a Gaussian velocity profile, rather than a constant speed. The peak velocity was 16.8 cm/s.

Each trial started by appearance of a small, white fixation point at the center of the visual display. The animals were required to maintain fixation within a 4 by 4 degree area around the fixation point for 150 ms to continue. Then optic flow appeared, simulating heading to the left or right relative to straight forward. In the early-target-onset paradigm, two choice targets would also be presented at the same time. Optic flow was presented for 1500 ms during which the animals were required to main central fixation. Then optic flow was turned off, and after another 50 ms, the central fixation point disappeared to indicate a go signal for choice towards one of the two choice targets. In the delayed-target-onset paradigm, the two choice targets would appear at this time. Correct choice led to reward of water.

### Electrophysiology

Electrophysiology data were collected from MT and MST in STS, and LIP in IPS. The ROIs were targeted based on the atlas, magnetic resonance imaging (MRI) scan, and electrophysiological characteristics. In STS, MT and MST were separated by receptive field properties. In particular, compared to MST, MT usually has a relatively smaller receptive field and located on the contralateral visual field. In IPS, a memory saccade task^59^ was used to localize LIP.

Spiking activity was collected by 16∼32-channel linear array (Plexon, and Clunbury Scientific LLC) and 4416-channel linear array (Neuropixels NHP 1.0, Belgium). Spike sorting was performed offline with the open-source Kilosort toolbox^60^. Subsequent unit selection was automated in Python based on predefined quality metrics, and recued with manual inspection. Specifically, units were retained if they met two criteria: (1) an average firing rate of at least 2 Hz, and (2) sustained spiking throughout the recording duration.

### Data analysis

#### Choice probability (CP)

CP was computed for each neuron in a way defined previously^2,8^. Briefly, neural firing rates across trials under identical stimulus condition were divided into two groups based on the animal’s choice. According to whether the choice was made consistent with or opposite to the neuron’s preferred motion direction, the two groups of data were defined as prefer-group and null-group, respectively. A receiver operating characteristic (ROC) curve was plotted^61^, with the horizontal axis representing the false alarm rate: proportion of trials within the null-group beyond the criteria, and the vertical axis representing the hit rate: proportion of trials within the prefer-group beyond the criteria. The area under the curve (AUC) was then calculated. If the neuron’s AUC value was greater than 0.5, it indicated that the neuron responded more strongly when the choice was made in the neuron’s preferred direction, and vice versa for area less than 0.5. CP not significantly different from 0.5 indicates a chance level: neural response is not correlated with the choice.

#### Targeted dimension reduction (TDR) and principle component analysis (PCA)

A model-based TDR (mTDR)^36^ was used, which is a linear regression-based model similar to TDR^62^. Specifically, it has explicit low rank regression parameter for task variable:

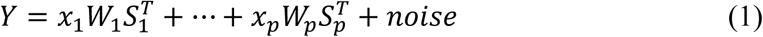

For our data, *Y* denotes the peristimulus time histogram (PSTH) of all neurons’ firing rate, a matrix with dimensions of number of units (U_n_) and time bins (T). The task related variable *p* included 6 variables: relative visual motion direction (ranged as: −2, −1, 0, 1, 2; represent moving direction sign × relative coherence), coherence (ranged as: 0, 1, 2; represent task difficulty from hard to easy), target color location (ranged as: −1, 1; represent red on left vs. green on left), perceptual choice (ranged as: −1, 1; choosing green vs. red), saccadic choice (ranged as: −1, 1; represent saccade left vs. right target), saccade in the last trial (ranged as: −1, 1; represent saccade left vs. right target). W_p_ is a weight matrix with size of U_n_ × *r*_*p*_, *r*_*p*_ is the subspace dimension number of variable *p*. *S*_*p*_ is a matrix with size of *T* × *r*_*p*_, representing the temporal component of each task-relevant variable *p*. *x*_*P*_ is a scalar indicating the value of the corresponding task variable for the current trial. We used the open-source toolbox^36^ which perform a marginal estimation to find subspace dimension *r*_*p*_, neural weighs *W*_*p*_, and basic function *S*_*p*_ for each variable.

After mTDR dimension reduction, each task related variable received a subspace. We then used sequential principle component analysis^36^ (seqPCA) to capture each subspace’s variance mainly explained axis along time. Specifically, population neurons were projected into a subspace and obtain an observation matrix 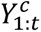 with size of *D* × *t*, where the superscript *c* denotes different trials. Data from all trials are concatenated along time *t* into a matrix 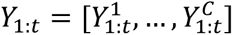 with size of *D* × *tC*. Then the explained variance of *Y*_1:*t*_ is given by:

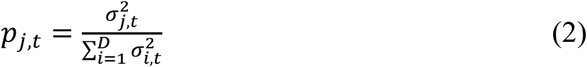

Within this subspace, when *σ*_1,*t*_ is the largest singular value, then *p*_1,*t*_ captures the maximum proportion of variance in the data over the early t time points. The value of *p*_*j*,*t*_ evolves over time depending on the structural characteristics of the data. If the variance of the data is entirely explained by the first principal component at all times, then *p*_1,*t*_ = 1 for all time points t. If the data lack structure, *p*_1,*t*_ will decrease monotonically over time until it converges to *p*_1,*t*_ = 1/*D*. If the data possess structure, *p*_1,*t*_ will increase linearly until a specific time point *t*^′^, after which it changes direction; this point *t*^′^ is referred to as the peak point. In this scenario, we compute the first singular vector of *Y*_1,*t*_′ up to time *t*^′^. Subsequently, we can examine the temporal evolution of the remaining variance of 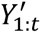 after removing the variance explained by the first singular vector.

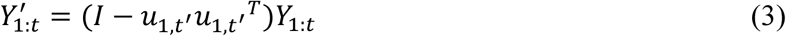

We then repeat the process above on 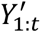 to find the second singular vector if it has further information.

### RNN simulation

#### Basic structure

Two modules were constructed in the network. The recurrent module 1 (128 units) received motion direction information input. Module 1 was further connected to the second module (256 units) through a feedforward projection. Information in the second module was read out by saccadic neurons for decisional choice. In addition, module 2 also received color information for perceptual choice. Module 2 sent feedback connection to module1. The network activity *V* follow a dynamical equation:

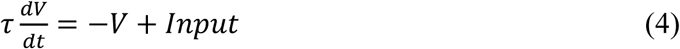

The *τ* is neural time constant, which is set to 20 ms for units in module 1 and 200 ms for units in module 2, to simulate relative faster and slow intrinsic dynamics in earlier sensory areas and higher level areas, respectively^63^. −*V* captures the intrinsic leaky process. In the discrete implementation, each time step was set to *dt* = 20 *ms*. The input for each recurrent module is defined as follows.

For input to module 1:

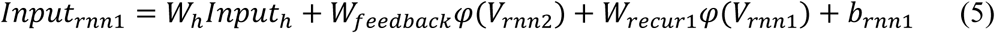

In the equation, *W*_*h*_*Input*_*h*_ represents the stimulus-driven input related to motion direction. *W*_*feedback*_*φ*(*V*_*rnn*2_) conveys feedback information from recurrent module 2 feedback. *W*_*recur*1_*φ*(*V*_*rnn*1_) captures the self-recurrent interactions within module 1. The constant bias *b*_*rnn*1_ accounts for the baseline activity of units in recurrent module 1.

For input to module 2:

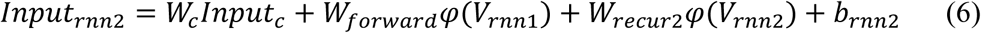

In the equation, *W*_*c*_*Input*_*c*_ represents the colored targets location information input delivered to module #2. The component *W*_*forward*_*φ*(*V*_*rnn*1_) conveys feedforward information from recurrent module 1. *W*_*recur*2_*φ*(*V*_*rnn*2_) reflects the self-recurrent interactions within module 2. The constant bias *b*_*rnn*2_ accounts for the baseline activity of units in module 2.

The activation function for units in the recurrent modules is *relu*:

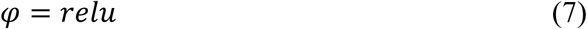

The readout of the network is defined as:

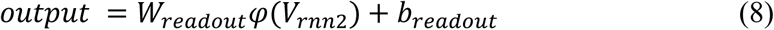

The *output* represents units activity modulated by saccadic direction, which is a linear readout of module 2. *W*_*readout*_denotes readout weight, and *b*_*readout*_ represents the baseline activity of output units.

#### RNN inputs

Our RNN input and output structure followed the design of previous studies^64^. One unit represented the fixation point, with its activity indicating fixation onset and offset. 32 units were tuned to motion direction in a range of 0 ∼2*π*. The activity amplitude *a*(*θ*) in response to the input of motion direction *θ* was dependent on its preferred motion direction *θ*_*i*_ defined as:

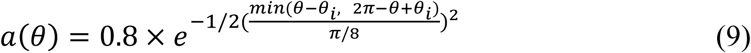

Motion directions with *θ* > *π* were assigned to T1 (red target), whereas directions with *θ* < *π* were assigned to T2 (green target).

To match neurophysiological experiments, we defined the temporal profile of heading stimulus as a Gaussian velocity waveform:

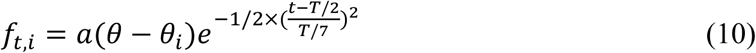

Thirty-two units represented the T1 target location, and the other 32 units represented the T2 target location. The input for the target location was defined as in equation (9). For each trial, T1 location was randomly selected from {0, *π*}, while the T2 location was set to be *π* relative to T1.

Noise was added to the network in two ways. The first type of noise was added to the motion direction input during the stimulus-on period. For each time point, the noise was generated as:

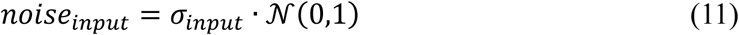

Where *N*(0,1) is a standard normal random variable and *σ*_*input*_ controls the noise amplitude. Noise was smoothed using a Gaussian filter with a kernel width of 7*σ*, where *σ* = 20 *ms*.

The second type of noise was added directly to the activity of units in recurrent modules 1 and 2, which was defined as:

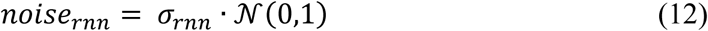

Where *σ*_*rnn*_ controls the amplitude of the recurrent noise and the term (*V*_*rnn*_ + 1) scales the noise relative to the current activity of each unit.

#### RNN outputs

32 units represented output neurons tuned to saccade direction. The expected activity of these neurons is defined in Equation (9).

The loss function was defined as the mean squared error between the actual outputs *y*_*it*_ and the expected output of *ŷ*_*it*_. A mask parameter *mask*_*it*_ was used to weight different time periods: The saccade-direction–encoding channels were weighted by 2 during the pre-response period, and by 10 during the response period. Formally, the loss was computed as:

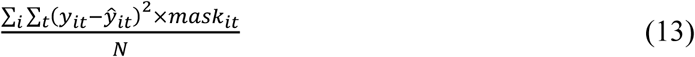

The network’s output saccade direction was computed as the vector sum of the activity of the 32 saccade-direction–tuned neurons. A trial was considered completed if the network’s output saccade direction was within 0.3*π* of either the T1 or T2 target location. For motion direction with *θ* > *π*, the correct saccade direction corresponded to T1 target location, whereas for *θ* < *π*, the correct saccade direction corresponded to the T2 target location.

#### Train batch

For each training batch, the motion direction was randomly selected from the range 0 to 2*π*. The coherence level for each batch was randomly chosen from {0.01,0.02,0.04,0.08}, and the stimulus duration was randomly selected from {400,800,1600} ms. Training was terminated once the network achieved a correct rate greater than 0.90.

#### Test batch

For trials in the test set, the motion direction was selected from ±0.2*π*. The coherence for each trial was chosen from {1,0.5,0.1,0.05,0}, and the stimulus duration was fixed at 1000 ms.

#### RNN neural activity

Neural activity was obtained from the test set. For each neuron, we calculated the perceptual choice preference and saccade choice preference based on the mean firing rate during the stimulus-on period. The choice-related divergence for each neuron was then computed as the difference between the normalized activity for the preferred direction and the non-preferred direction. The population-level choice-related divergence signal was obtained by averaging the choice-related divergence across all units. Statistical significance was assessed using a t-test comparing the observed activity to a shuffled condition activity (p < 0.001).

## Acknowledgements

We thank Wenyao Chen, Qi Zhao for monkey care and training, and Ying Liu for C++ software programming. This work was supported by grants from the Strategic Priority Research Program of the Chinese Academy of Sciences (XDB1010000) to Y.G.

## Author Contributions

N.Z. and Y.G. conceived the project and designed the experiments. N.Z. performed the experiments, constructed RNN simulations, and analyzed the data. N.Z. and Y.G. wrote the manuscript.

