## supplemental figure for "Context-Dependent Motor Feedback Underlies Choice-Related Signals in Visual Cortex"

### Supporting information

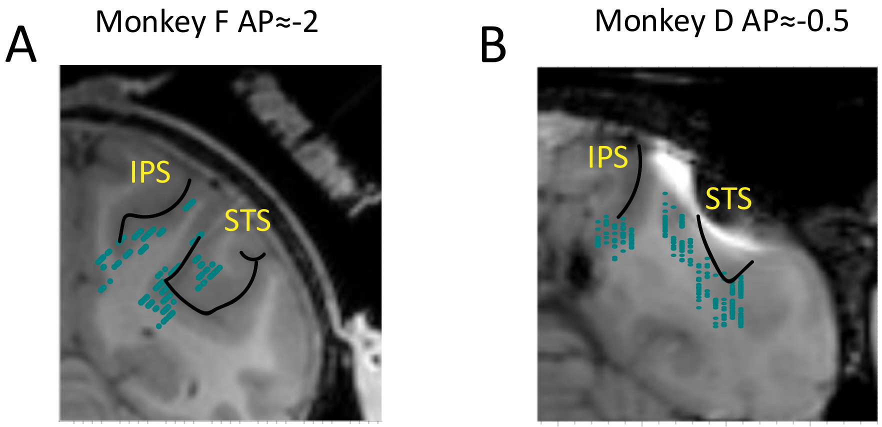


**Figure S1. Recording sites of MT, MST in STS, and LIP in IPS**

(A) Coronal section of MRI image of monkey F around AP = -2 mm. All the recording sites between AP -2.6 ~ -1.3 mm were compressed into a single section as shown in the figure, leading to data that were seemingly out of ROI.

(B) Similar format in (A), but for monkey D in the section of AP = -0.5 mm.

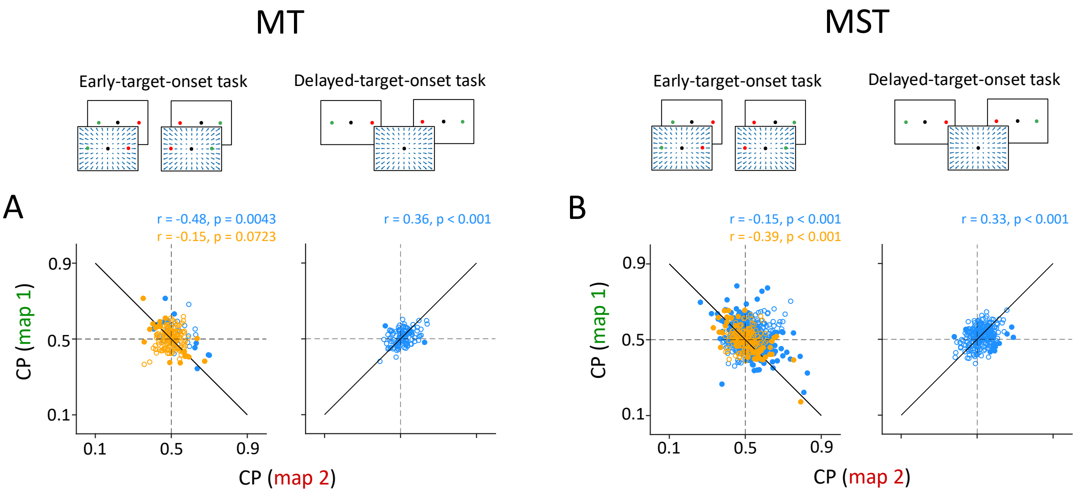


**Figure S2. CP results separated for area of MT and MST**

(A) Comparison of individual neuron's CP under the two-map conditions in MT. CP measured under the early-target-onset and delayed-target-onset tasks are on the left and right in each plot, respectively. Filled and open dots represent neurons with significant and insignificant saccadic choice component, respectively. Embedded texts inform Pearson correlation of CP measured between the two maps for each animal (blue, monkey F; orange, monkey D). Solid line is unity line. Horizontal and vertical dashed lines are chance level of CP (0.5).

(B) Similar format in (A), but for area MST.

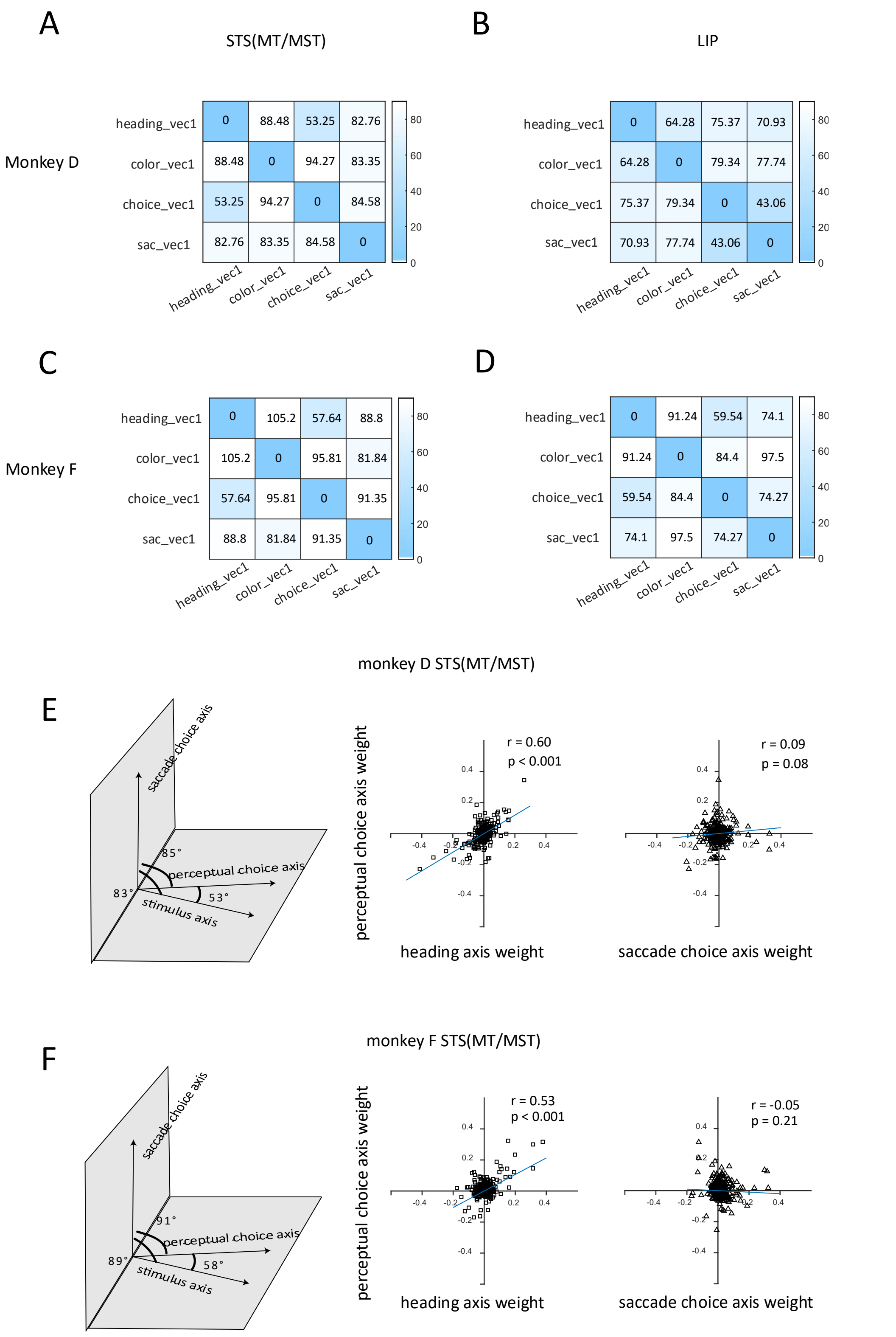


**Figure S3. Relationship between stimulus, perceptual choice, and saccadic choice axis extracted from the mTDR & seqPCA dimension reduction**

(A-D) Relationship between sequential axis in the two monkeys and two areas. Each value in the heat map is the angle between two axes. Higher luminance indicates larger angle difference. Vec1: the first sequential vector.

(E-F) Left: Relative angle of the sensory, perceptual choice and saccadic choice axis in the two animals (top vs. bottom). Middle: Comparison of each neuron's weight for sequential #1 heading axis and sequential #1 perceptual choice axis. Right: Comparison of each neuron's weight for sequential #1 saccadic choice axis and sequential #1 perceptual choice axis. Cyan line in the scatter plot is a reference line with slope equal to correlation coefficient r (Pearson correlation).

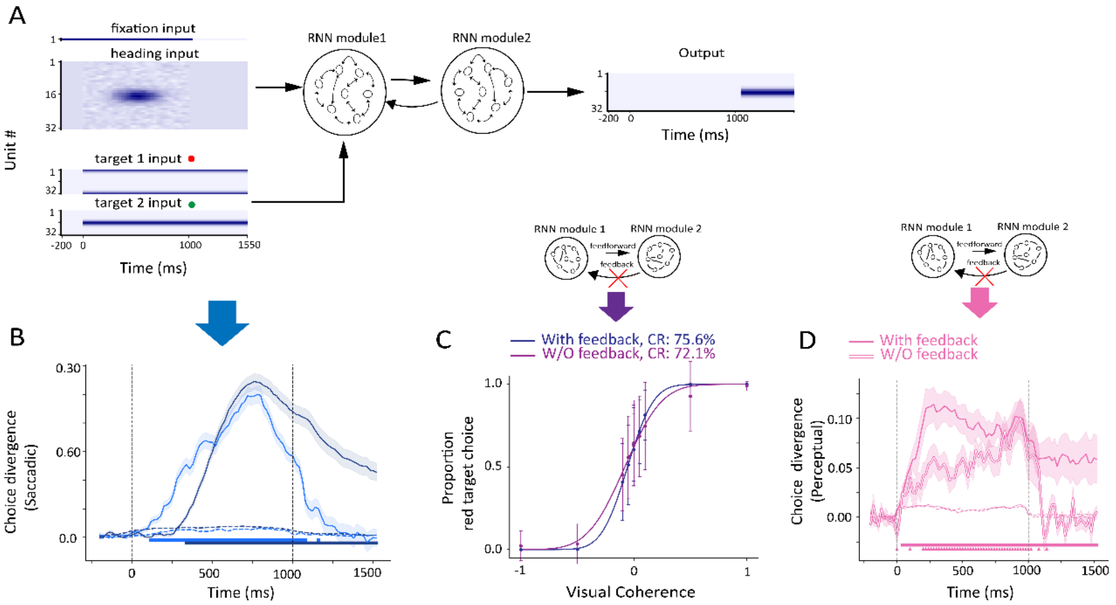


**Figure S4. RNN manipulations**

(A) Structure of two-module RNN with colored-targets wired into module #1 instead of module #2.

(B) The saccadic choice signal in module #1 (light blue) was leading that in module #2 (dark blue) of an example network. Shaded area represents error bar of 1 SEM.

(C) Behavioral performance when removing feedback connections from module #2 to module #1. The blue line is the average behavior performance of 16 networks, error bar represents 1SD. The magenta line is the average behavior performance of the same 16 networks but with removed feedback connection.

(D) Same as the module #1 data in the Figure 5C but with manipulation of the RNN feedback connections from module 2 to module 1. Squares and triangles represent significant perceptual signals before and after removing the feedback, respectively.
